# A loss-of-function variant in GFRAL associates with increased alcohol consumption in humans

**DOI:** 10.64898/2026.03.06.709997

**Authors:** Johanne Marie Justesen, Kimmie Vestergaard Sørensen, Jayashri Seshadri, Jens Skjoldan Svenningsen, Matthew Aguirre, Yosuke Tanigawa, Robert Minneker, Amalie Rasmussen Lanng, Cecilie Bæch-Laursen, Emilie Skytte Andersen, Louise J. Skov, Sebastian B. Jørgensen, Ulrik Becker, Filip Krag Knop, Manuel Rivas, Niels Grarup, Matthew Paul Gillum

## Abstract

Alcohol is an ancient and enduring component of the human diet, yet it is a dose-dependent cytotoxin and teratogen, raising the possibility that endogenous, state-dependent mechanisms constrain intake. Growth differentiation factor 15 (GDF15) is an endocrine hormone that rises during pregnancy—predominantly via secretion from blastocyst-derived placental trophoblasts into the maternal circulation—and is also induced in other tissues, particularly hepatocytes, by toxins and cellular stress. However, its function in humans remains unclear. Here, we show that circulating GDF15 levels are elevated 5-fold in individuals with alcohol dependence, identify a rare loss-of-function variant in the GDF15 receptor gene GFRAL associated with approximately 2.6 additional UK alcohol units (∼21 g ethanol) per week, and demonstrate that recombinant GDF15 reduces alcohol drinking in mice. Collectively, these findings support a model in which GDF15 acts as an endocrine signal induced by chronic alcohol exposure—and potentially during pregnancy—to limit alcohol intake in humans.

**Highlights:**

- GDF15 is markedly elevated in humans with alcohol dependence
- A truncating GFRAL variant associates with higher alcohol intake in UK Biobank
- GFRAL frameshift disrupts GDF15–RET signaling in vitro
- Recombinant GDF15 suppresses voluntary alcohol intake in mice

**Graphical abstract:** 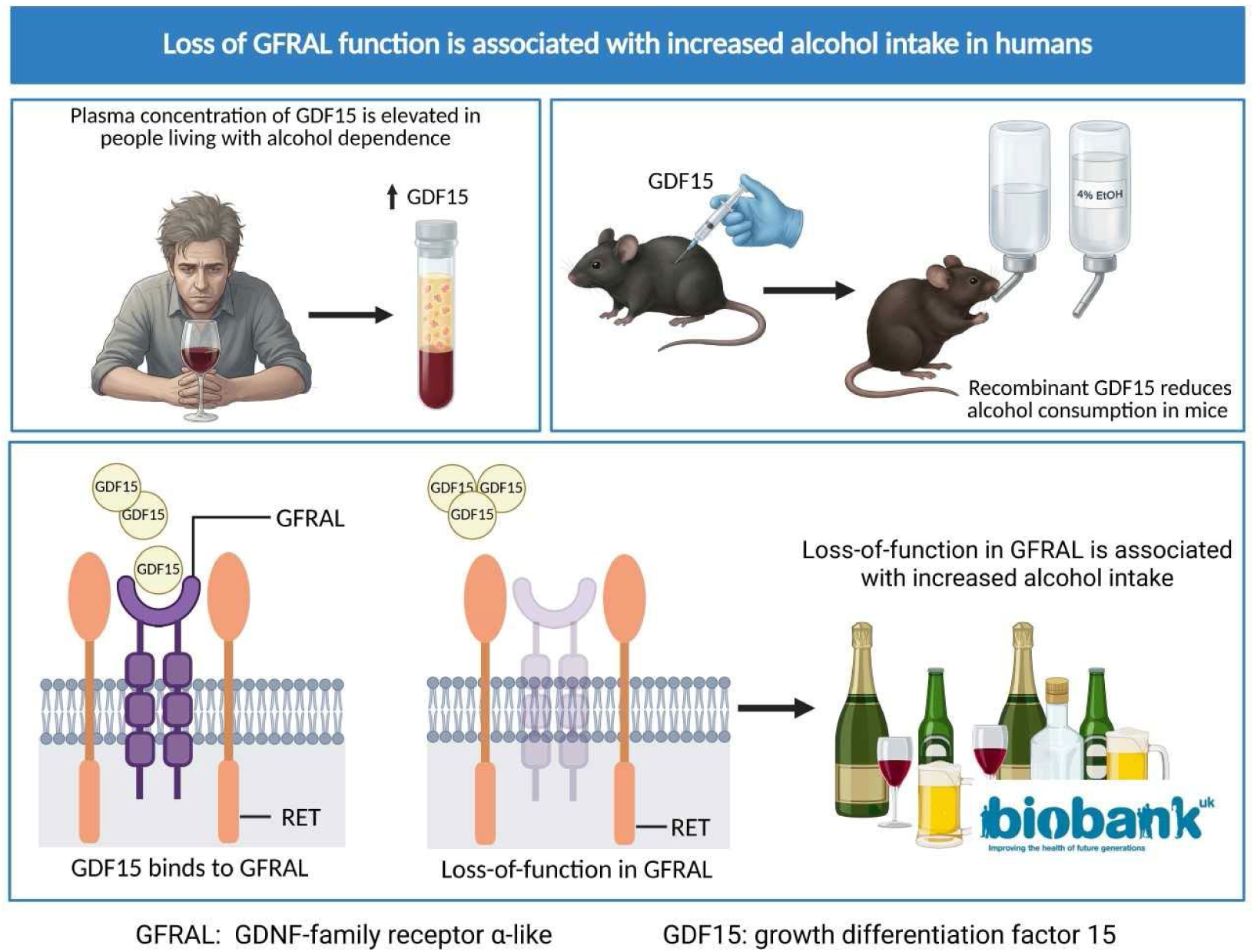

## Introduction

Growth differentiation factor 15 (GDF15) is a unique circulating member of the transforming growth factor β superfamily that was identified in the 1990s as a macrophage-derived cytokine (initially termed macrophage inhibitory cytokine-1) [1]. In rodents, GDF15 overexpression or administration reduces food intake and body weight [2–4]. In humans, circulating GDF15 increases in several diseases such as cancer and also physiologically during pregnancy. Circulating GDF15 levels correlate positively with obesity but remain stable in response to changes in nutrient availability, metabolic hormones, or body weight [5–11]. It is thus unclear why GDF15 should inhibit food intake in model organisms, since appetite regulators are usually induced postprandially (e.g., glucagon-like peptide 1) or are sensitive to changes in fat mass (e.g., leptin) [12–14]. Moreover, a clinical trial with the long-acting GDF15 receptor agonist LY3463251 reported negligible weight loss over 12 weeks in humans, suggesting functions for GDF15 beyond regulating overall energy balance and body weight [15].

GDF15 binds a receptor complex in the brainstem composed of GFRAL and the receptor tyrosine kinase RET [16–19]. The expression of GFRAL is limited to the area postrema, a brain area involved in nausea and food aversion, leading to the theory that GDF15 is a “non-homeostatic” (or “allostatic”) regulator of appetite, secreted in response to ingested toxins may not be required for the maintenance of normal energy balance [16–19]. Supporting this idea, cytotoxic chemotherapeutics cause GDF15-dependent anorexia in mice, and GFRAL-RET signaling activates neurons in the central nucleus of the amygdala via the parabrachial nucleus, which is an aversive neural network also engaged by other toxic insults such as lithium chloride and lipopolysaccharide [20]. Therefore, production of GDF15 by injured cells may be a general mechanism by which chemically heterogeneous cytotoxins are sensed, their ongoing consumption halted, and their future consumption avoided.

Alcohol is a significant source of energy in the Western diet constituting 5-10% of total energy intake [21] yet can dose-dependently damage multiple organ systems and the developing fetus [22], making it a potential target for regulation by GDF15. Moreover, under physiological conditions—apart from pregnancy—GDF15 is primarily liver-derived and upregulated under some of the same conditions as fibroblast growth factor 21 (FGF21), a hepatic stress hormone whose sequence variants—and variants in its co-receptor *KLB*—are associated with alcohol consumption and misuse in humans [23–25]. FGF21 selectively inhibits alcohol and sweet appetite and increases 40-fold acutely after alcohol drinking in humans [25–27]. Given the co-secretion of FGF21 and GDF15 by stressed cells, and the fact that alcohol increases GDF15 production *in vitro* and in rodents [28], we hypothesized that GDF15 might be induced by alcohol drinking to subsequently reduce the desire to drink alcoholic beverages.

In this study, we examine the relationship between alcohol consumption and the GDF15-GFRAL system in humans and mice to understand if it might be involved in controlling alcohol intake.

## Results

### GDF15 secretion is not regulated by acute alcohol intake in humans

Because FGF21 levels are rapidly increased by alcohol drinking, we first investigated whether GDF15 secretion was acutely induced in 12 individuals who, as part of the AlcoGut study, received ethanol equivalent to five standard drinks (60 g) via a nasogastric tube, reaching a peak plasma alcohol concentration of 1.7 ± 0.3 g/L (SEM) [29]. In contrast to FGF21, circulating GDF15 levels decreased non-significantly to a nadir at 90 min after dosing before returning to baseline over the remainder of the 4 h observation period (**Figure 1A**).

**Figure 1.**
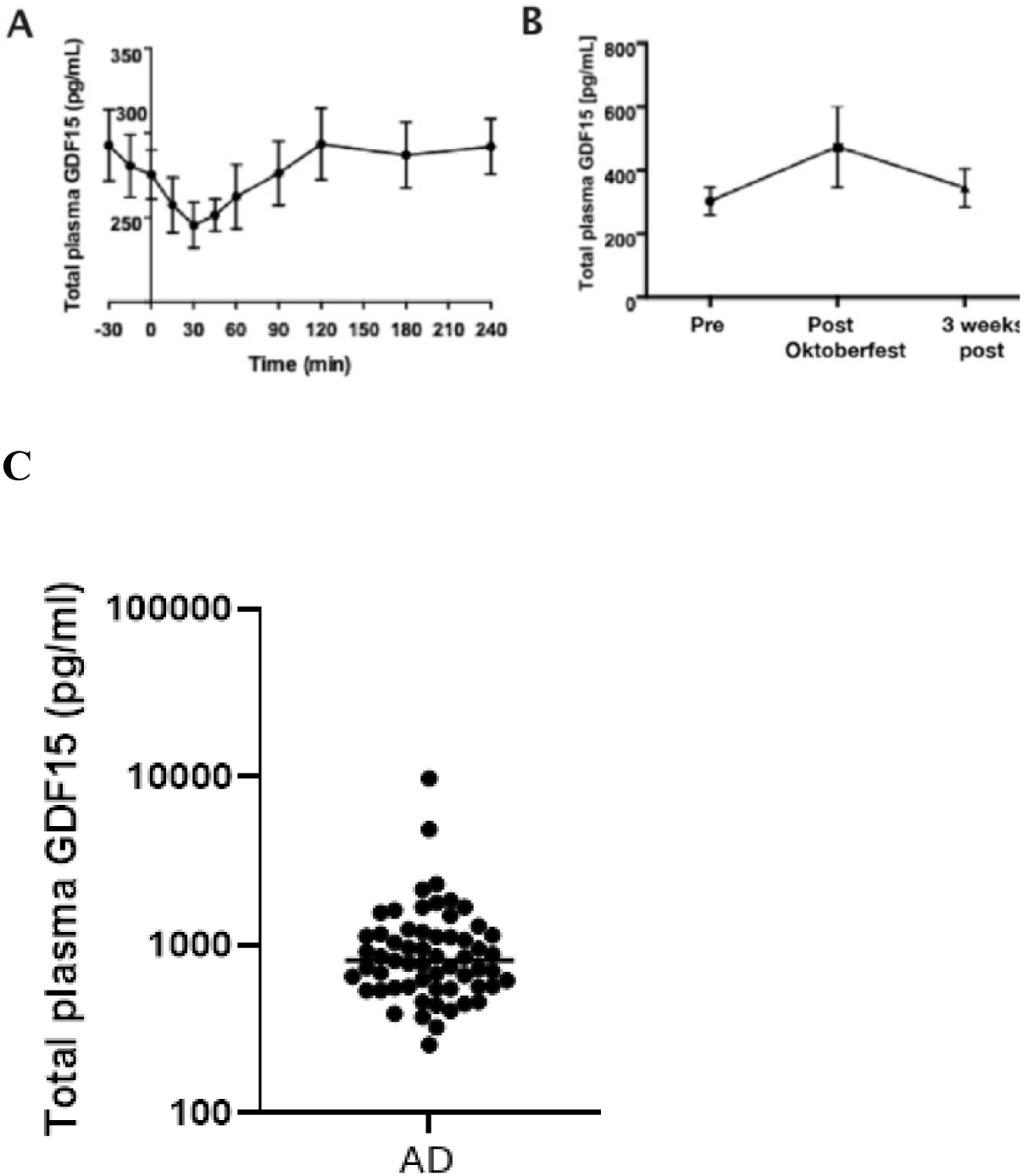
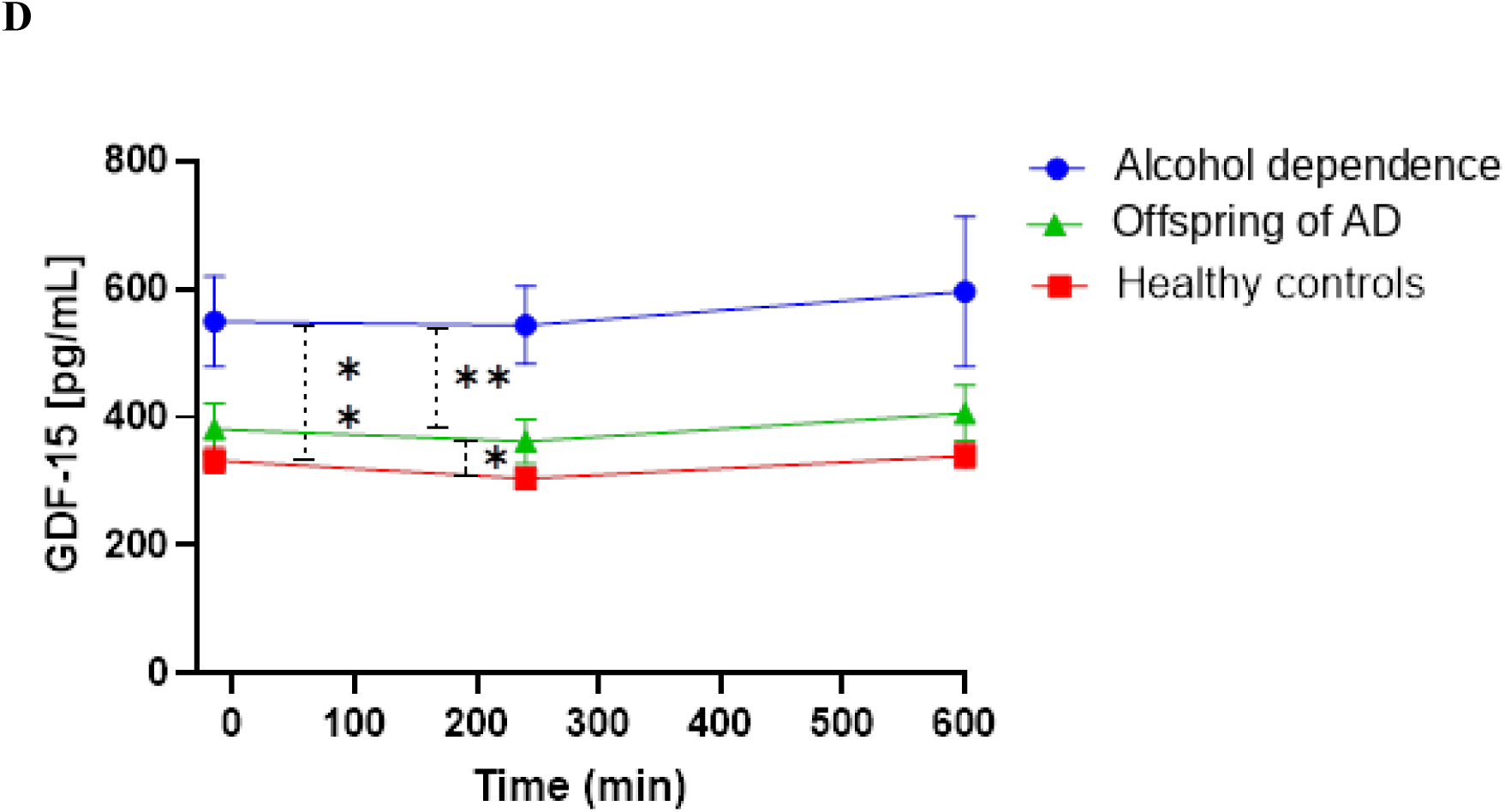
Circulating GDF15 levels and alcohol consumption. A) Plasma GDF15 levels from -30 to 240 minutes in young healthy humans (N = 12) who received a single intragastric bolus of alcohol equivalent to the amount found in five standard drinks (60 g). B) GDF15 plasma levels in three healthy human males before, 32 hours after, and three weeks after consuming an average of 22 beers/day for three consecutive days during Oktoberfest. C) Plasma GDF15 levels in patients seeking outpatient treatment for alcohol dependence (N=59). D) GDF15 plasma levels in participants with alcohol dependence (N=15), a family history of alcohol dependence (N=15), or controls (N=15) given an oral bolus of 0.5 g/kg at t=0. Data are presented as mean ± SEM and differences were tested by repeated measures analysis of variance (A, B, and D), with Tukey’s multiple comparisons test to evaluate group differences (D).

### GDF15 levels tend to increase after a weekend of binge drinking in humans

Next, we addressed whether circulating GDF15 levels would be affected by a period of heavy drinking for three days at Oktoberfest. We measured GDF15 in fasting blood samples collected from three healthy volunteers before, immediately after, and three weeks after attending the three-day festival [26]. During Oktoberfest, individuals consumed an average of 22 beers per day for three consecutive days. Although the sample was small (N = 3), limiting statistical inference, plasma GDF15 tended to increase immediately after the festival relative to before but reverted to baseline levels after three weeks of follow up (**Figure 1B**).

### GDF15 levels are markedly elevated in patients with alcohol dependence and increased in the offspring of males with alcohol dependence

The observation that GDF15 levels tended to increase in response to sub-chronic, but not acute, alcohol intake in humans led us to measure GDF15 levels in 59 outpatient, treatment-seeking individuals with alcohol dependence (median age 49 years) and generally normal liver function [30]. Circulating GDF15 increases with age, and levels in middle-aged Danish individuals (40–64 years) are 198 ± 88 (SD) pg/mL in females and 224 ± 107 (SD) pg/mL in males [31]. In contrast, plasma GDF15 levels in patients with AUD ranged from 257 to 9,814 pg/mL, with a mean of 1,132 ± 1,342 (SD) pg/mL (**Figure 1C**).

To investigate this phenomenon further in a highly controlled setting, we measured circulating GDF15 levels in 15 male participants with alcohol dependence, 15 healthy male offspring of fathers with alcohol dependence, and 15 healthy males with no family history of alcohol dependence, both at baseline and in response to an oral bolus of alcohol (0.5 g/kg). Participants were matched for age, body weight, height, BMI, fasting glucose, hemoglobin, ALT, AST, bilirubin, INR, cholesterol, and transient elastography (dB and kPa) [32]. As expected, participants with AUD reported significantly higher daily alcohol consumption (7.6 units/day vs. 1 in predisposed individuals and 0.5 in controls), higher Alcohol Use Disorders Identification Test (AUDIT) scores (22 vs. 7.8 and 5.5), and higher gamma-glutamyl transferase (47 U/L vs. 20.1 and 21.4 U/L). GDF15 levels did not increase acutely in any group in response to alcohol intake. However, baseline GDF15 levels were markedly elevated in participants with AUD (563 ± 17 (SEM) pg/mL) and slightly elevated in the offspring of males with AUD (384 ± 13 (SEM) pg/mL) relative to controls (326 ± 11 (SEM) pg/mL) (Figure 1D).

### Human carriers of a rare protein-truncating variant in *GFRAL* consume more alcohol

Protein-coding genetic variants that are predicted to result in loss of protein function by either introducing a stop codon, structural changes by frameshift or disruption of a splice site, provide direct biological insight [33]. Thus, we searched for protein-truncating variants (PTVs) with a minor allele count of more than 200 (resulting in minor allele frequency (MAF) > 0.00059) in GDF15, GFRAL and RET in unrelated white British individuals from the UK Biobank population study (N = 337,122) to study their associations with weekly total alcohol consumption. Only one PTV was identified by this approach: a frameshift variant (rs527905870, consequence: p.Ile339AsnfsTer36) with MAF of 0.00096 (MAF comparable to gnomAD database, Supplementary Table 1) in the C-terminal region of GFRAL. This frameshift of GFRAL protein structure at amino acid position 339 is predicted to result in a GFRAL receptor lacking the entire transmembrane and cytoplasmic domains (Supplementary Figure 1). This variant was associated with an increased weekly total alcohol intake in carriers compared to non-carriers (beta 0.12 SD, p = 0.009, Table 1). Thus, carriers of GFRAL p.Ile339Asnfs consume ∼2.6 additional UK units/week (∼21 g ethanol) compared with non-carriers.

**Table 1.** Association of protein truncating variants in GFRAL with weekly alcohol intake in 337,119 white British individuals from UK Biobank

| Gene | Variant | MAF<sup>a</sup> | Consequence | Beta<br>(SD<sup>b</sup>) | SE<sup>c</sup> | p-value |
| --- | --- | --- | --- | --- | --- | --- |
| *GFRAL* | rs527905870 | 0.00096 | p.Ile339AsnfsTer36<br>(frameshift) | 0.12 | 0.04 | 0.009 |

<sup>a</sup>Minor allele frequency, MAF
<sup>b</sup>Standard deviation, SD
<sup>c</sup>Standard error, SE

### *GFRAL* p.Ile339Asnfs did not associate with adverse phenotypes

A phenome-wide association study (PheWAS) of the *GFRAL* PTV can potentially predict adverse effects of therapeutically targeting this receptor complex and pathway. Hence, this approach can offer insights into potential beneficial effects as well as the adverse drug effects, in addition to providing supportive details regarding plausible associations with alcohol intake and GFRAL biology in general. We did not observe associations with phenotypes indicating severe effects from partial inhibition of GFRAL (**Supplementary Table 2**). However, we do not observe individuals homozygous for *GFRAL* PTVs in the UK Biobank, and cannot exclude that such individuals could potentially have severe consequences including lethal effects. Nevertheless, it was recently reported that complete GDF15 loss in humans does not lead to an overt phenotype, suggesting that this scenario is unlikely if GFRAL primarily functions as a receptor for GDF15 [34].

### *In vitro* characterization shows that GFRAL function and glycosylation of its co-receptor RET is disrupted by the p.Ile339Asnfs variant

Next, we sought to characterize the functional consequences of the GFRAL p.Ile339Asnfs variant in vitro. HEK293T cells co-transfected with RET and the p.Ile339Asnfs GFRAL variant failed to induce AKT phosphorylation in response to GDF15, in contrast to cells expressing wild-type GFRAL (**Figure 2A**), consistent with prior reports that GDF15–GFRAL signaling requires RET to elicit intracellular signaling [16–19]. We also observed a strikingly different RET gel migration pattern between cells expressing wild-type GFRAL versus the variant receptor, with variant-transfected cells lacking the 170 kDa RET band (**Figure 2B**). The nascent 120 kDa RET protein undergoes two sets of post-translational glycosylations during maturation, first giving rise to a 150 kDa protein found in the endoplasmic reticulum, and then to the fully mature 170 kDa protein, which resides in the cell membrane [36,37]. To test whether this difference reflected differential glycosylation, samples from Figure 2B were treated with PNGase F to remove all N-linked oligosaccharides (**Figure 2C**). PNGase F treatment yielded a single 120 kDa band irrespective of expressed GFRAL receptor, suggesting that wild-type, but not variant, GFRAL expression increases the abundance of mature 170 kDa RET that mediates intracellular GDF15 signal transduction [19].

**Figure 2.**
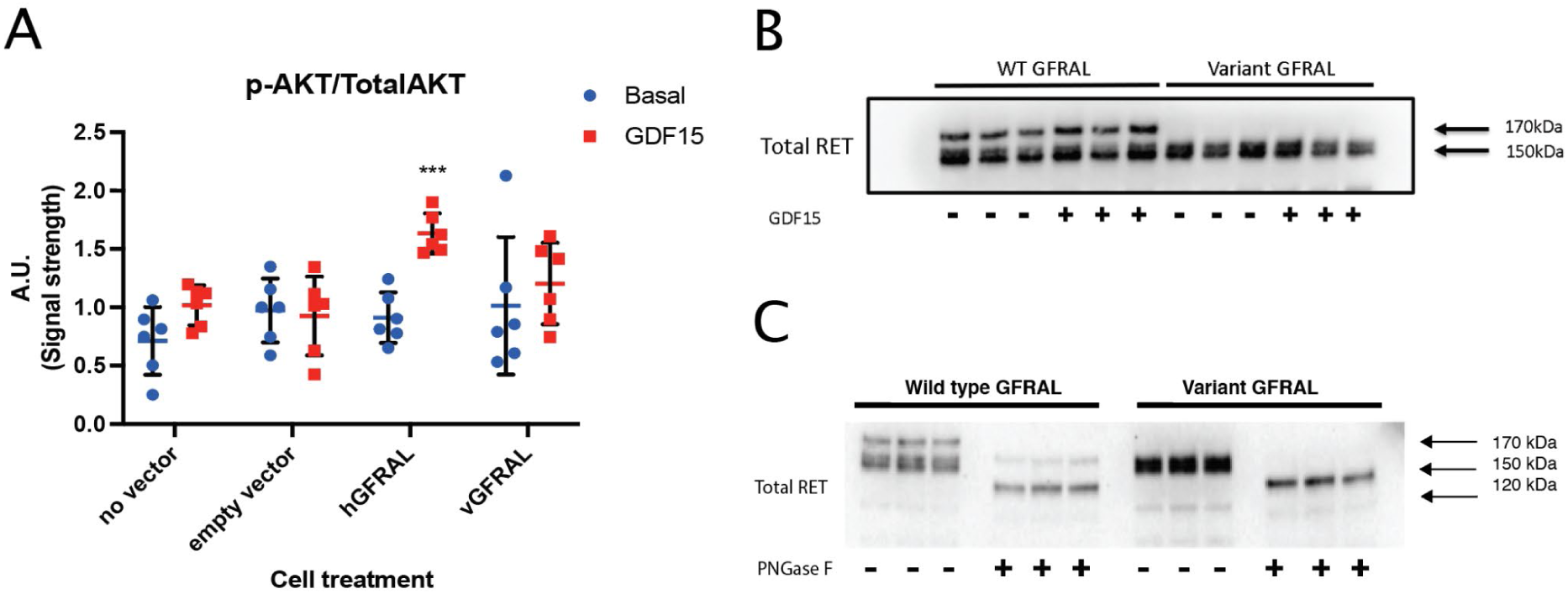
In vitro characterization shows that GFRAL receptor function is disrupted by p.Ile339Asnfs variant. A) GDF15 signal transduction characterized in samples with no vector, empty vector, wild-type GFRAL vector (hGFRAL) and p.Ile339Asnfs GFRAL vector (vGFRAL), blotted for total and phosphorylated AKT. B) Differential RET maturation (glycosylation) depends upon the presence of receptor in the cell membrane. C) PNGase F assay of the wild-type GFRAL and the p.Ile339Asnfs GFRAL. Data are presented as mean ± SEM, with each group compared to its corresponding empty vector control by unpaired two-sided t-test.

### Recombinant human GDF15 reduces alcohol intake in mice

Finally, we tested whether GDF15 could suppress alcohol intake in ad libitum chow-fed mice also provided with a choice between a 4% (v/v) ethanol bottle and a water bottle. In this two-bottle choice preference paradigm, single 1 mg/kg subcutaneous injection of recombinant human GDF15 acutely reduced 24 h alcohol drinking by 60% compared to vehicle-treated mice (**Figure 3**). By comparison, chow intake was reduced 38% by GDF15 (2.1 ± 0.47 g/mouse vs. 3.4 ± 0.46 g/mouse, p<0.0001).

**Figure 3.**
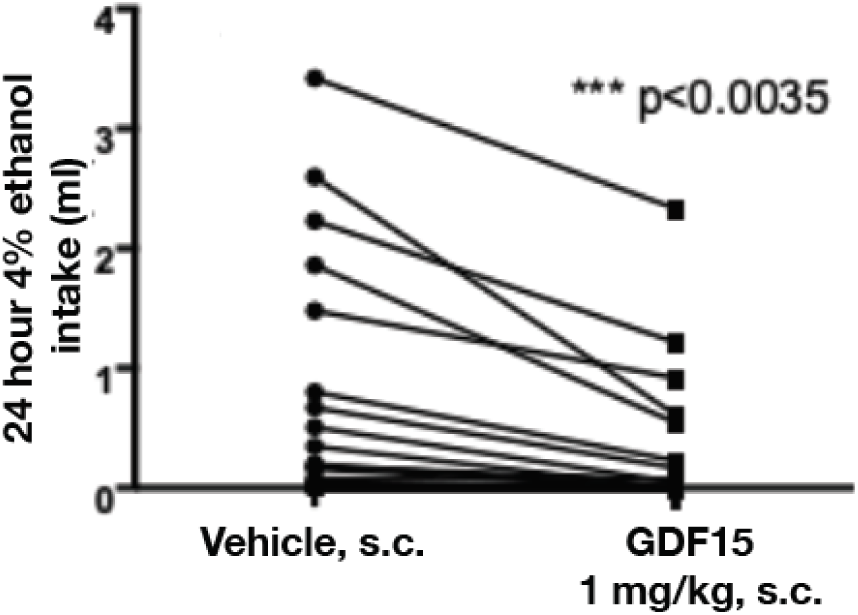
Recombinant human GDF15 reduces alcohol intake in mice. The effect of 1 mg/kg rhGDF15 on 24 h alcohol intake in mice (N = 30). Data are presented as mean ± SEM, and the significance of the treatment was determined by paired t-test.

## Discussion

The cytotoxic and teratogenic effects of dietary alcohol represent evolutionary pressures that may have favored biological mechanisms that constrain intake. Analogous to negative-feedback systems that regulate feeding, such constraints could include endocrine signals that report alcohol exposure or alcohol-induced cellular stress to the brain. Although alcohol intake is shaped by many central neurobiological pathways (including GABAergic, glutamatergic, opioidergic, and others), endocrine factors provide an additional, systemic layer of feedback that can couple peripheral ethanol metabolism to drinking behavior. In this context, FGF21 is one example of a liver-derived hormone that is induced by alcohol and has been implicated genetically and experimentally in limiting alcohol consumption, including in non-human primates [38,39]. Here, we show that GDF15 displays features consistent with a longer-timescale endocrine regulator of alcohol intake: circulating GDF15 is increased in humans living with alcohol dependence to levels only otherwise seen in cancer and pregnancy [40], GDF15 is induced by chronic (but not acute) alcohol intake, a rare loss-of-function variant in its receptor (GFRAL) is associated with increased alcohol consumption, and recombinant GDF15 preferentially reduces voluntary alcohol drinking in mice.

The tissue sources of GDF15 are also consistent with a role in linking physiological state to alcohol-related behaviors. During pregnancy, GDF15 is predominantly secreted into maternal circulation by placental trophoblasts derived from the developing conceptus, whereas outside pregnancy it is produced by diverse cell types, particularly hepatocytes, in response to toxins and cellular stress. These features are compatible with the idea that GDF15 serves as a systemic signal of physiological vulnerability to cause protective changes in behavior, and they motivate future studies testing whether GDF15 contributes to the altered alcohol preferences and alcohol aversion frequently reported during human pregnancy [41].

It has been proposed that GDF15 functions more broadly to limit intake of cytotoxic ingested substances, acting as a negative-feedback signal generated in response to cellular stress and sensed in the brainstem via binding to GFRAL in the area postrema [42–45]. Notably, however, GDF15 was not acutely increased by alcohol in our data, arguing against a simple model in which alcohol triggers the canonical “acute toxin” GDF15 response. One alternative is that sustained elevation of GDF15 during chronic heavy drinking modulates neural processing of alcohol reward, reinforcement, or preference rather than mediating an acute aversive response. This interpretation is consistent with prior evidence that GDF15 can influence food preferences and macronutrient selection [46]. A key question raised by our human findings is how elevated GDF15 in alcohol dependence relates to ongoing heavy drinking: one possibility is that chronic GDF15 induction reflects a compensatory, but insufficient, feedback signal that is overridden by the neuroadaptations and learned motivational processes characteristic of dependence. Another non-mutually exclusive possibility is that sensitivity to GDF15’s central effects could be diminished in established alcohol dependence (e.g., functional “resistance” at the level of downstream circuitry), such that elevated circulating levels do not necessarily translate into proportionately increased behavioral restraint. Discriminating among these possibilities will require longitudinal human studies and mechanistic work in model systems that better capture dependence-associated neuroadaptations.

The genetic association between the inactivating protein-truncating variant in GFRAL (p.Ile339Asnfs) and higher alcohol intake in the UK Biobank supports a physiological role for the GDF15–GFRAL pathway in regulating drinking behavior at the population level, thus primarily at typical, non-pathological levels of drinking. Importantly, in our PheWAS analysis this variant did not associate with body mass index, suggesting that its association with alcohol intake is not simply secondary to energy-intake or obesity-related phenotypes that have previously been linked to the GDF15/GFRAL locus [47–49]. In addition, rare loss-of-function variation in GFRAL has been reported to associate with bilirubin in GeneBass, consistent with the regulation of serum bilirubin levels by alcohol exposure [50] and the idea that GFRAL loss-of-function increases alcohol intake, which in turn contributes to altered bilirubin; however, direct effects of the pathway on liver-related traits or independent pleiotropic effects cannot be excluded.

Our observation that RET is essential for GDF15–GFRAL signaling via AKT is consistent with prior reports [16–19], but, to our knowledge, the finding that GFRAL expression alters the abundance of the doubly glycosylated mature form of its co-receptor RET has not been described previously. One potential mechanism is that functional GFRAL at the cell surface promotes stabilization or recycling of mature RET to the plasma membrane. These observations motivate further studies to clarify how GFRAL regulates mature RET abundance and glycosylation state, and how this phenomenon shapes signaling dynamics downstream of GDF15.

This study has several limitations. First, our human alcohol-consumption measures are based on self-report, which introduces measurement error; however, self-reported alcohol intake can show reasonable agreement with collateral and other external sources, and there are few feasible alternatives at scale. Second, residual confounding is a key concern in observational comparisons involving cohorts with alcohol use disorder, although some evidence suggests that patients with dependence may resemble the general population more closely than often assumed [30]. Accordingly, we cannot conclude with certainty that the differences in circulating GDF15 observed in alcohol dependence are exclusively attributable to alcohol exposure itself, as comorbidities, medication use, diet, and liver disease severity may also contribute. Finally, there are notable species differences in the pharmacological effects of GDF15, including far more pronounced body-weight effects in rodents than in humans, and this limitation may extend to alcohol-related behavioral effects as well [15].

Together, our data support a model in which GDF15 is induced by alcohol exposure and can act through GFRAL to constrain alcohol drinking. These findings motivate further work to determine whether augmenting this pathway has therapeutic value in alcohol use disorder or in alcohol-related liver disease. At the same time, because individuals with alcohol dependence exhibit markedly elevated endogenous GDF15, any translational strategy will need to address whether (and in whom) additional pharmacological activation of the pathway provides benefit beyond an already engaged endogenous response, including the possibility of partial pathway resistance at the level of downstream circuitry. Nevertheless, elevated endogenous hormone levels do not preclude pharmacological efficacy, because therapeutics can achieve supraphysiological exposures that overcome reduced sensitivity; a conceptual analogy is type 2 diabetes, where insulin resistance is overcome by increasing exogenous insulin dosing to achieve glycaemic control. Accordingly, therapeutic utility may depend on disease stage, patient subgroups, or the ability of long-acting agonists to engage central GFRAL signaling more effectively than endogenous GDF15.

## Acknowledgements

The Novo Nordisk Foundation Center for Basic Metabolic Research is an independent Research Center at the University of Copenhagen partially funded by an unrestricted donation from the Novo Nordisk Foundation (Grant number: NNF18CC0034900, http://cbmr.ku.dk/). This research has been conducted using the UK Biobank Resource (http://ukbiobank.ac.uk/). We thank all the participants in the UK Biobank study. The primary and processed data used to generate the analyses presented here are available in the UK Biobank access management system (https://amsportal.ukbiobank.ac.uk/) for application 24983, “Generating effective therapeutic hypotheses from genomic and hospital linkage data” (http://www.ukbiobank.ac.uk/wp-content/uploads/2017/06/24983-Dr-Manuel-Rivas.pdf), and the results are displayed in the Global Biobank Engine (https://biobankengine.stanford.edu). We also thank the volunteers participating in the clinical studies from which plasma samples were analyzed for GDF15.

J.M.J. was supported by grant NNF17OC0025806 from the Novo Nordisk Foundation and the Stanford Bio-X Program. M.A.R. is supported by Stanford University and a National Institute of Health center for Multi- and Trans-ethnic Mapping of Mendelian and Complex Diseases grant (5U01 HG009080).

## Author Contributions

Conceptualization: J.M.J., K.V.S., L.J.S., S.B.J., F.K.K., M.R., N.G., M.P.G.

Methodology: J.M.J., J.S., J.S.S., M.P.G.

Investigation: J.S., J.S.S., M.P.G.; M.P., A.R.L., E.S.A., F.K.K., U.B.

Data curation: J.M.J., M.A., Y.T., M.R.

Formal analysis: J.M.J., M.A., R.M., M.R., N.G.

Writing – original draft: J.M.J., K.V.S., N.G., M.P.G.

Writing – review & editing: all authors. Supervision: N.G., M.R., F.K.K., M.P.G.

## Materials and Methods

### Lead contact

Further information and requests may be directed to and will be fulfilled by the lead contact, Matthew Gillum

### AlcoGut study

The study was approved by the Scientific-Ethical Committee of the Capital Region of Denmark (identification no. H-16026085) and the Danish Data Protection Agency and registered on ClinicalTrials.gov (identifier: NCT03348371). Primary results of the study have previously been reported [29], and the present GDF15 analyses were performed post hoc. The study was conducted according to the Declaration of Helsinki and written informed consent was obtained from all participants. Twelve healthy Caucasian men were included (age 20–50 years; BMI 19–25 kg/m²; weekly alcohol intake <14 Danish units, where 1 unit = 12 g ethanol). Key exclusion criteria included first-degree relatives with type 1 diabetes, type 2 diabetes or liver disease; diagnosed liver disease; alcohol-related disease; and nephropathy. The study consisted of two experimental days performed in randomized order. In a double-blind fashion, alcohol was administered intragastrically through a nasogastric tube or intravenously aiming at isoethanolaemia, respectively. Participants abstained from alcohol for 5 days before each visit and met after an overnight fast (10 h). An intravenous catheter was inserted into an antecubital vein in each arm (one for infusion and one for blood sampling), and a nasogastric tube was inserted. The hand and forearm of the blood-sampling arm were wrapped in a heating pad (∼42°C) throughout the experiment for arterialization of the blood. At time 0 min, participants received alcohol (0.70 g/kg; 20% solution) mixed in isotonic saline water via the gastric tube over 5 min, or the same dose infused intravenously over 45 min, as described previously [29]. Blood samples were collected at -30, -15, 0, 15, 30, 45, 60, 90, 120, 180, and 240 min. All tubes were centrifuged for 15 min at 2,000 g and 4°C, and plasma/serum were stored at -20°C and -80°C, respectively, until analyses.

### Oktoberfest study

Three healthy Danish men (aged 42 years, BMI of 23.8, 21.4, and 28.3 kg/m², respectively) with moderate, social drinking habits contacted our clinic for a health examination before three days of Oktoberfest participation in Munich, Germany. Upon their return they requested a reevaluation of their metabolic health (32 hours after consuming the last unit of alcohol at Oktoberfest), as well as three weeks later. All agreed to have their cases reported anonymously. Subjects were examined in the morning after an overnight (10 h) fast. Body weight, height, body composition (by bioimpedance), and hepatic stiffness (by fibro scanning) were evaluated, and blood and urine were sampled [26].

### Individuals with alcohol dependence

Patients with alcohol dependence from the Greater Copenhagen area who were attending an outpatient clinic at Hvidovre University Hospital, Denmark, were consecutively enrolled over a 9-month period as originally described [30]. Treatment at hospitals and hospital outpatient clinics is free in Denmark and admission is open. Seventy percent of included patients were men; the median age was 49 years (interquartile range (IQR) 43–54 years); the median duration of addiction was 14 years (IQR 6–21 years); the median alcohol intake in the months before admission was 171 g/day (IQR 86–288 g/day); and the median BMI was 25 kg/m² (IQR 22–29 kg/m²). Patients fulfilling the ICD-10 diagnostic criteria for alcohol dependence and seeking assistance at the clinic were asked to participate. As part of the clinical evaluation and planning of treatment, we used the Addiction Severity Index, from which composite scores for the alcohol profile were calculated [51]. All participants provided informed consent, and the study was performed according to the Declaration of Helsinki. The Danish National Committee on Biomedical Research Ethics (ref. no. H-A-2007-0032) and the Danish Data Protection Agency (ref. no. 2007-41-0735) approved the study.

### Measurement of GDF15 levels and their response to acute alcohol ingestion in participants with alcohol dependence, the offspring of patients with alcohol use disorder, and healthy controls

GDF15 levels in plasma were measured in samples from study NCT03892369, whose primary endpoint was the between-group difference in plasma FGF21 response to alcohol ingestion evaluated by area under the curve (AUC) [32]. Total human GDF15 was measured using the Quantikine ELISA kit (R&D Systems, DGD150). The study was approved by the Scientific Ethical Committee of the Capital Region of Denmark (no. H-18063495) and conducted according to the Declaration of Helsinki. Included were 15 male participants with alcohol dependence (ICD-10 code F10.2), 15 healthy male participants with a father diagnosed with AUD (predisposed), and 15 healthy male participants without any family history of AUD (controls). Key inclusion criteria were men aged 20–65 years and BMI 19–27 kg/m². Participants were matched for age, body weight, height, BMI, fasting glucose, hemoglobin, ALT, AST, bilirubin, INR, cholesterol, and transient elastography (dB and kPa). Participants met after an overnight fast, were placed in a semi-recumbent position, and an intravenous catheter was placed for blood sampling (with heating for arterialization). At time 0 min, participants consumed alcohol (0.5 g/kg; 37.5% solution [v/w]; 15.5 kJ/kg) over 10 min. Blood samples were drawn and VAS questionnaires were completed hourly for 10 h (until 600 min). After 180 min, participants received a standardized meal (spaghetti Bolognese).

### GDF15 and acute alcohol preference study in mice

Thirty-two naïve female C57BL/6 mice (10 weeks old) were implanted with subcutaneous radiofrequency identification (RFID) chips and housed in groups of four in an automated HM2 rodent feeding system. Mice were kept in a temperature- and humidity-controlled specific-pathogen-free facility under a 12:12 h light–dark cycle. The HM2 system records drinking data from two channels with single-mouse resolution based on RFID detection when the animal enters the drinking chamber, and each bottle is mounted on a scale to detect spillage. Because individual mice showed stable day-to-day alcohol intake (range: 0–4 mL/24 h), each mouse served as its own control on successive test days. Two animals were excluded due to RFID chip loss or failure. Mice were acclimated with free choice between water and 4% (v/v) ethanol in water for 7 days. After intake stabilized, mice were treated immediately prior to the start of the dark phase with vehicle or GDF15 (1 mg/kg), and fluid intake was recorded for the subsequent 24 h.

### Human Genetic studies

Biobank is a large-scale population-based cohort study with the overall aim to improve prevention, diagnosis, and treatment of a wide range of serious and life-threatening illnesses. In 2006-2010, the UK Biobank recruited 502,650 participants aged 37-73 years at 21 centers across England, Wales and Scotland using standardized procedures [52]. When participants agreed to take part in UK Biobank, they visited the closest assessment center for collection of baseline information, physical measures, and biological samples. They have undergone measures, provided blood, urine, and saliva samples for future analysis, provided detailed information about themselves and agreed to have their health followed. The UK Biobank protocol is described in detail online (http://www.ukbiobank.ac.uk/wp-content/uploads/2011/11/UK-Biobank-Protocol.pdf).

The participants have been extensively examined. Outcome events are captured via registries, and there is an expert adjudication group who reviews medical records of every register-reported case of specified conditions.

### Genotyping and quality control of the UK Biobank sample

Genotyping and imputation procedures for the UK Biobank dataset have been previously described [52]. Briefly, two genotyping arrays, the UK Biobank Axiom Array (N = 438,427) and the UK BiLEVE Axiom Array (N = 49,950), were used to create the final genotype release of 805,426 loci for 488,377 individuals. Genotype quality control was performed before the data were released publicly, including removing participants with excess heterozygosity or missingness rate, and removing markers showing effects related to batch, plate, sex, or array, or those demonstrating discordance across control replicates. We searched for protein truncating variants in GDF15, GFRAL and RET with a minor allele count of at least 200. This cutoff will result in a MAF > 0.00059 (Supplementary Table 1).

### Alcohol phenotype definition

Participants were asked to report their average weekly alcohol consumption through a self-reported touchscreen questionnaire. We calculated the weekly alcohol unit intake by combining intake questions of beer/cider, red wine, white wine/champagne, spirits, and fortified wine and multiplied by the standard drink size and alcohol content. Given responses for each individual (N = 348,583) who took part in the alcohol section of the survey and lifestyle questionnaire, we performed the following steps: 1) For each type of beverage, impute uninformative non-missing values (e.g. “I don’t know” or “Prefer not to answer”) with the median consumption rate among informative responses; 2) Sum self-reported consumption of all beverages, weighted by estimated alcohol content (https://www.drinkaware.co.uk/alcohol-facts/alcoholic-drinks-units/what-is-an-alcohol-unit/), to get net weekly alcohol consumption; 3) Exclude individuals who self-report consuming more than 250 units of alcohol per week — this is the same cutoff used by UK Biobank to reject survey responses as invalid; 4) Replace self-reported values of zero units per week by one unit per week, as all individuals participating in the survey indicated that they consume alcohol; and 5) Apply population-level rank-transformation to the resulting sum of alcohol content. The resulting distribution of alcohol consumption is extremely heavy tailed, with mean weekly consumption estimated to be 23.5 units (8g pure alcohol; SD 19.9 units) among alcohol drinkers.

## Data availability

The UK Biobank data is available through the UK Biobank (http://ukbiobank.ac.uk/). GWAS results are available on the Global Biobank Engine (https://biobankengine.stanford.edu). Further requests for data are available upon correspondence with the Lead Contact.

## Statistical analyses

After quality control we identified one PTV in *GFRAL* having MAF > 0.0005 and this variant was examined in relation to weekly alcohol consumption in the UK Biobank using PLINK v2.00a (2 April 2019). We performed linear regression (via the plink2 --glm option) for the variant, within a subset of unrelated white British individuals (N = 337,122; 242,297 with self-reported alcohol consumption data). We apply quantile normalization to the phenotype “weekly alcohol intake”, as well as the raw derived values. Association analysis was adjusted for age, sex, array, and four population-genetic principal components.

We performed a PheWAS of the GFRAL rs527905870 using results from GWAS that has previously been described by [53] and present results from traits that were associated with p < 1×10^-4^ and for disease traits having at least 100 cases.

Since the variant is rare, we examined the genotyping plot using Scattershot (http://mccarthy.well.ox.ac.uk/static/software/scattershot/) to exclude genotyping errors (Supplementary Figure 2).

## Cellular signaling characterization of rs527905870

Functional signaling assay. We performed functional characterization assays to investigate whether the predicted protein truncation affects the PTV’s ability to relay signal to downstream components in the GDF15–GFRAL axis. An in vitro model was set up using HEK293 cells transiently transfected with bicistronic vectors expressing the co-receptor RET along with either wild-type GFRAL or the rs527905870 variant. Cells expressing GFRAL and RET were serum-starved for 4 h in serum-free media containing 1% penicillin/streptomycin and then treated with 100 nM GDF15 for 15 min. After stimulation, cells were lysed in RIPA buffer and sonicated. Samples were analyzed by immunoblotting. Primary antibodies against total ERK1/2, phospho-ERK1/2, total AKT, phospho-AKT, total RET, and GAPDH were purchased from Cell Signaling Technology. Band intensities were quantified using ImageJ, and statistical analyses were performed in GraphPad Prism using an unpaired two-tailed t-test.

Deglycosylation assay. To examine the basis for the differential RET migration pattern, we performed deglycosylation using PNGase F (New England Biolabs). HEK293 cells were transfected with bicistronic vectors expressing RET along with either wild-type GFRAL or the PTV. Twenty-four hours after transfection, cells were lysed in RIPA buffer and sonicated. Samples were treated with PNGase F and prepared for western blotting according to the manufacturer’s instructions. For immunoblotting, an antibody against total RET (Cell Signaling Technology) was used.

## Supplemental Information Titles and Legends

**Supplementary Table 1:** Population frequencies for GFRAL rs527905870 from gnomAD.

| Population | Allele count | Allele number | Allele frequency |
| --- | --- | --- | --- |
| African | 7 | 24752 | 0.0002828 |
| Ashkenazi Jewish | 46 | 10328 | 0.004454 |
| East Asian | 0 | 19816 | 0.000 |
| European (Finnish) | 0 | 25068 | 0.000 |
| **European (non-Finnish)** | **182** | **128426** | **0.001417** |
| Latino | 33 | 35180 | 0.0009380 |
| South Asian | 5 | 30348 | 0.0001648 |
| Other | 10 | 7170 | 0.001395 |
| Female | 137 | 128510 | 0.001066 |
| Male | 146 | 152578 | 0.0009569 |
| **Total** | **283** | **281088** | **0.001007** |

Population frequency for *GFRAL* rs527905870 in UK Biobank (minor allele frequency (MAF)
= 0.00096) is similar to gnomAD European population (MAF = 0.0014).
https://gnomad.broadinstitute.org/variant/6-55264036-C-CA?dataset=gnomad_r2_1

**Supplementary Table 2:** Results from a Phenome-wide association study (PheWAS) of GFRAL rs527905870 in white British individuals from UK Biobank (p < 1×10^-4^).

| Phenotype name | Cases | Controls | OR/Beta | SE | *p*-value |
| --- | --- | --- | --- | --- | --- |
| Fracture thumb | 424 | 336965 | 1.98 | 0.414 | 1.63e-06 |
| Gastrointestinal bleeding | 134 | 336965 | 2.47 | 0.587 | 2.52e-05 |
| Ease of skin tanning (Never tan, only burn) | 59404 | 330847 | 0.348 | 0.095 | 2.61e-04 |
| Eye infection | 358 | 336965 | 1.775 | 0.505 | 4.34e-04 |
| Crumbed or deep-fried poultry intake (Diet, 24hr recall, Online follow-up) | 9169 | 9167 | -0.543 | 0.159 | 6.63e-04 |
| Bile duct obstruction / Ascending cholangitis | 1312 | 336965 | 1.180 | 0.357 | 9.60e-04 |

Analyses including diseases/traits with more than 100 cases. OR, odds ratio; SE, standard error.
Only phenotypes with $p < 1 \times 10^{-4}$ are presented.

## Supplementary figures

**Supplementary Figure 1:**
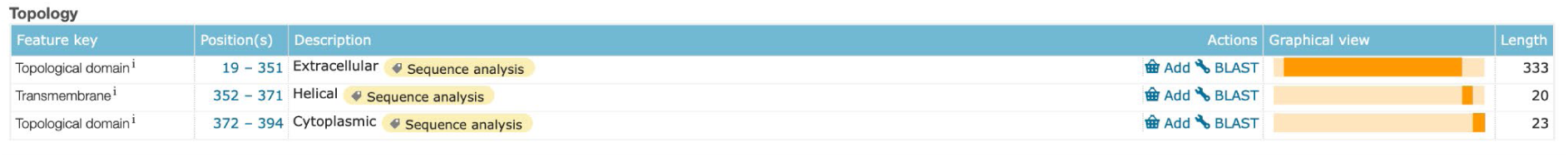
Topology information on GFRAL from UniProt Database. Uniprot topology information https://www.uniprot.org/uniprot/Q6UXV0. The rs527905870 results in frameshift of GFRAL protein structure at amino acid position 339 (p.Ile339AsnfsTer36) resulting in a GFRAL receptor possibly lacking the transmembrane and cytoplasmic parts of the receptor.

**Supplementary Figure 2:**
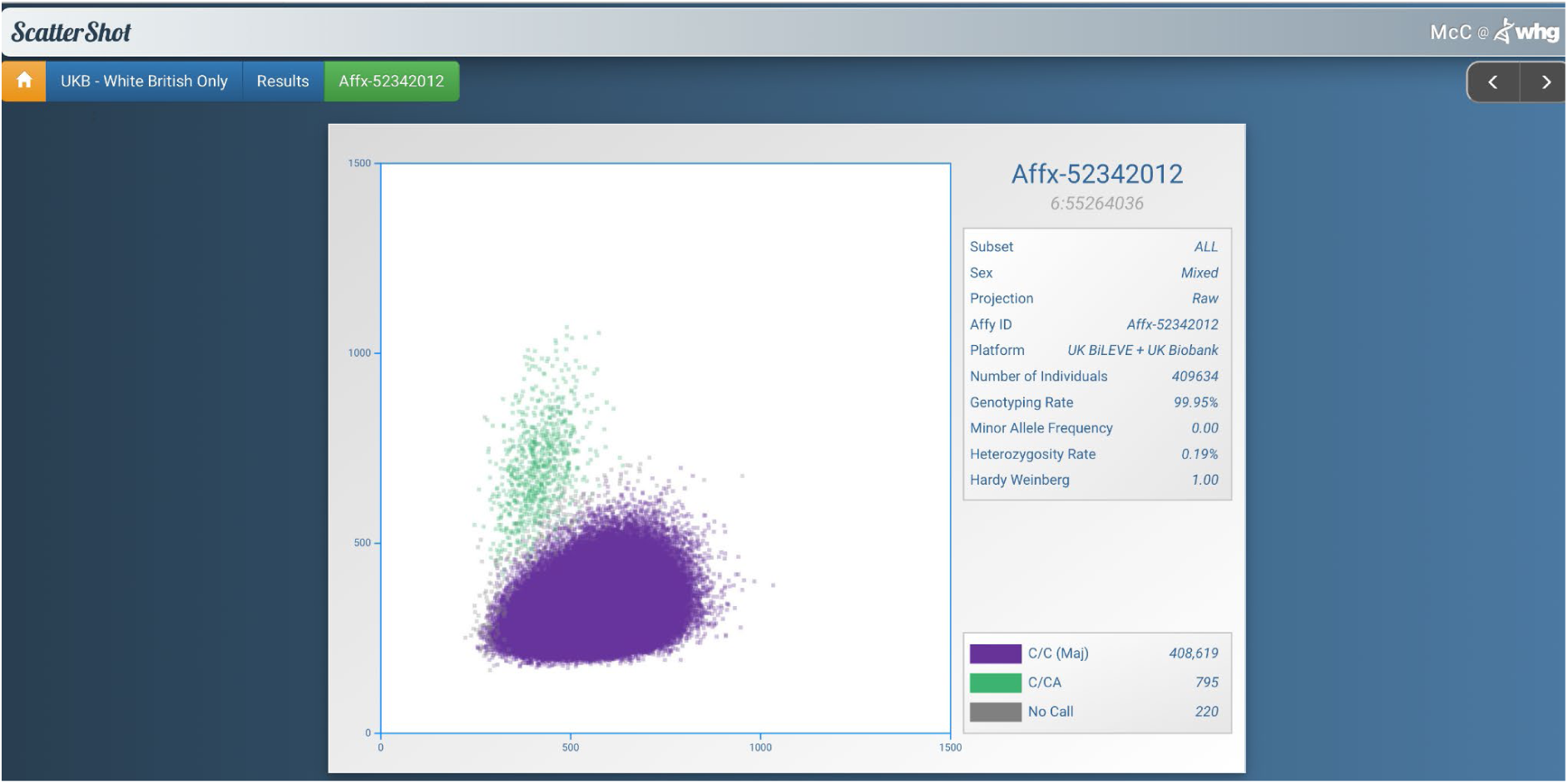
Scattershot showing genotype calling for GFRAL rs527905870 variant in UK Biobank http://mccarthy.well.ox.ac.uk/static/software/scattershot/ Genotype calling for the rs527905870 in white British individuals from UK Biobank (N = 409634) (Affx-52342012). There are 795 of the heterozygous genotype C/CA of rs527905870 among White British individuals.

